# Transient Vascular Occlusions in a Zebrafish Model of Tuberculous Meningitis

**DOI:** 10.1101/2025.02.28.640927

**Authors:** Megan I. Hayes, Sumedha Ravishankar, Victor Nizet, Cressida A. Madigan

**Affiliations:** School of Biological Sciences, University of California, San Diego; La Jolla, 92037, United States of America; Department of Pediatrics, University of California, San Diego; La Jolla, 92037, United States of America

## Abstract

Tuberculous meningitis (TBM), caused by *Mycobacterium tuberculosis,* is a severe manifestation of tuberculosis that occurs when the bacteria invade the brain. In addition to extensive inflammation, vascular complications such as stroke frequently arise, significantly increasing the risk of disability and death. However, the mechanisms underlying these vascular complications remain poorly understood, as current knowledge is derived exclusively from human studies. To date, no animal model has been established to investigate the onset and progression of vascular pathology in TBM. Here, we use transparent zebrafish larvae to investigate vascular pathology during the early stages of mycobacterial brain infection, establishing a model for studying TBM-associated vascular complications. We find that mycobacteria preferentially attach to the lumen of vessel bifurcations and induce vessel enlargement. These attached microcolonies are sufficient to occlude brain blood vessels in the absence of an organized thrombus. The majority of microcolony-associated occlusions are transient and contribute to global hypoperfusion of the brain. These vascular disruptions lead to accumulation of oxidative stress and cell death in both the vasculature and neurons. Taken together, these findings demonstrate the occurrence of ischemic events during the early stages of mycobacterial brain infection and establish an animal model for studying vascular complications in TBM.

## Introduction

Tuberculous meningitis (TBM), the deadliest manifestation of *M. tuberculosis* infection, claimed the lives of 78,300 children and adults in 2019 ^1-3^. Cerebrovascular complications are particularly common in TBM and are associated with increased mortality ^4,5^. These complications typically manifest as vasculitis and/or stroke and are strong predictors of disability and death in TBM ^6^.

TBM is thought to occur when *M. tuberculosis* disseminates beyond the initial infection site in the lungs and travels through the bloodstream to the brain ^2^. Though still debated, various studies in the last several decades have begun unraveling the mechanism by which mycobacteria enter the brain parenchyma ^7-9^. Once in the brain, *M. tuberculosis* infection is associated with the formation of myeloid cell aggregates, known eponymously as Rich foci, as well as vascular pathologies such as vasculitis and stroke ^4,10^. While these findings have expanded our understanding of how mycobacteria enter the brain and initiate Rich foci formation, the mechanisms underlying TBM-associated vascular pathologies remain incompletely understood.

Much of our current knowledge regarding TBM’s impact on the brain’s vasculature derives from human magnetic resonance imaging (MRI), computed tomography (CT), and autopsy studies ^5,11^. These studies have provided critical insights into the clinical presentation of TBM-associated vascular complications. However, findings are inconsistent across human studies. This is likely due to the underreporting of TBM, variable access to treatment, and the often-silent nature of vascular complications ^2,5,12^. Furthermore, the observational nature of these human studies limits our ability to establish causal relationships between vascular pathology and TBM.

To elucidate the cellular and molecular mechanisms underlying TBM-associated vascular complications, animal models are essential. Zebrafish larvae have been established as a useful model for studying mycobacterial brain infections ^8,13^. In this model, larvae are infected with a pathogenic relative of *M. tuberculosis, M. marinum*, which replicates many features of tuberculosis infection is zebrafish larvae ^8,13^. In addition to the extensive genetic tools available in zebrafish, their optical transparency at the larval stages allows for real-time visualization of mycobacterial brain infection in a live host ^14,15^. This model presents an exciting opportunity to investigate the vascular complications of TBM in vivo. Using live confocal imaging, we characterized the early vascular consequences of mycobacterial brain infection and found that microcolonies are associated with altered vascular morphology and transient occlusion of brain blood vessels. These disruptions contribute to a global reduction in cerebral blood flow velocity, as well as oxidative stress and cell death in both the vasculature and neurons. Our findings lay the groundwork for future investigations into TBM-associated vascular pathology using the zebrafish model.

## Materials and Methods

### Zebrafish husbandry and infections

Zebrafish husbandry and experiments were conducted in compliance with guidelines from the U.S. National Institutes of Health and approved by the University of California San Diego Institutional Animal Care and Use Committee and the Institutional Biosafety Committee of the University of California San Diego. Wildtype AB strain zebrafish or transgenics in the AB background were used, including Tg(*fliE:GFP*) ^16^, Tg(*flk:GFP*) ^17^, Tg(*gata:DsRed*) ^18^, Tg(*moesin:GFP*) ^19^, Tg(*cd41:GFP)* ^20^, and Tg(*nbt:dsred*) ^21^. Larvae were anesthetized with 2.8% Syncaine (Syndel #886-86-2) prior to imaging or infection. Larvae of indeterminate sex were infected by injection of 10 nL into the caudal vein at 3 days post fertilization (dpf) using a capillary needle containing bacteria diluted in PBS + 2% phenol red (Sigma #P3532), as previously described ^22^. Titered, single-cell suspensions were prepared for all *M. marinum* strains prior to infection by passing cell pellets from mid-log phase cultures (OD600 = 0.5 ± 0.1) repeatedly through a syringe to remove clumps, as described ^22^. After caudal vein injections, the same needle was used to inject onto 7H10 (Sigma-Aldrich # M199) agar plates containing 50 μg/mL hygromycin B (ThermoFisher #10687010) or 50 μg/ml kanamycin (TCI #K0047) in triplicate to determine colony forming units (CFUs) in the inoculum. ∼100-500 CFUs of wildtype *M. marinum* were administered to the larvae for experiments unless otherwise specified. After infection, larvae were housed at 28.5°C, in zebrafish water containing ddH^2^O, 14.6 g/l sodium chloride (JT Baker #3628-F7), 0.63 g/l potassium chloride (Sigma-Aldrich #P3911), 1.83 g/l calcium chloride (G-Biosciences #RC-030), 1.99 g/l magnesium sulfate heptahydrate (MP Biomedicals #194833), methylene blue chloride (Millipore Sigma #284), and 0.003% 1-phenyl-2-thiourea (PTU, Sigma-Aldrich #189235) to prevent melanocyte development.

### Bacterial strains

*M. marinum* M strain (ATCC #BAA-535) tdTomato or eBFP under control of the msp12 promoter ^22,23^, were grown in 50 μg/mL hygromycin B (ThermoFisher, #10687010) or 50 μg/mL kanamycin (TCI, # K0047) in liquid culture, consisting of 7H9 Middlebrook medium (Sigma-Aldrich, #M0178) supplemented with 2.5% oleic acid (Sigma, #O1008), 50% glucose, and 20% Tween-80 (Sigma, #P1754) ^22^. Agar plates contained 7H10 Middlebrook agar (HiMedia, #M199) supplemented with oleic acid, albumin (Sigma, #A9647), dextrose, and Tween-80 ^22^.

### Dextran and stain administration to zebrafish larvae

To visualize vessels in larvae without transgenic fluorescent vessels and to test luminal perfusion, 10 kDa Alexa647 Dextran (ThermoFisher, #D22914) was diluted to 1 mg/ml in PBS and injected into the caudal vein at the time of imaging. To visualize oxidative stress, cellROX (ThermoFisher, Cat# C10422) was diluted to 25 μM and injected into the hindbrain ventricle immediately prior to imaging. To visualize cell death, SYTOX deep red nucleic acid stain (ThermoFisher, Cat# S11380) was diluted to 10 μM and injected into the hindbrain ventricle.

### Zebrafish larva microscopy and image analysis

For confocal imaging, larvae were embedded in 1.2% low melting-point agarose (IBI Scientific #IB70051) ^22^. A series of z stack images with a 0.82-1 µm step size were generated through the brain using the Zeiss LSM 880 laser scanning microscope with an LD C-Apochromat 40x objective. Imaris (Bitplane Scientific Software) was used to measure fluorescence intensity and construct three-dimensional surface renderings ^24^. When events were compared between larvae or in paired vessels, identical confocal laser settings, software settings, and Imaris surface-rendering algorithms were used. For imaging blood vessels, transgenic animals with fluorescent blood vessels Tg(*fliE:GFP*) ^16^ or Tg(*flk:GFP*) ^17^, were used, or Alexa647 Dextran (ThermoFisher, #D22914) was injected intravenously.

For RBC tracking, larvae were embedded in 1.2% low melting-point agarose (IBI Scientific #IB70051) ^22^. Fast frame image was performed on a Leica SP8 resonant scanning laser confocal microscope with a 25x water objective. Images were acquired at a frame rate of 83.3 fps for a total of 10.02 seconds. To track cells, the Imaris spot algorithm was used and applied at all time points to track individual RBCs.

### Experimental reproducibility and statistical analysis

Many experiments were repeated multiple times to ensure reproducibility. The number of experimental replicates is indicated in the corresponding figure legend. If no number is listed, the experiment was conducted once. The following statistical analyses were performed using Prism 8 (GraphPad): Student’s and paired t-test, and chi-squared test, and one-way ANOVA with multiple comparisons. The statistical tests used for each figure can be found in the corresponding figure legend. The *n* values for larvae and microcolonies are given below each corresponding graph.

## Results

### Mycobacteria attach to brain microvascular bifurcations and alter vessel morphology

Attachment of microcolonies to brain microvascular endothelial cells is thought to be one of the earliest steps in mycobacterial invasion of the brain ^7^. In many neurovascular diseases, specific vessels or regions are disproportionately impacted. For instance, atherosclerosis, stroke, and aneurysms often occur at arterial bifurcations ^25,26^. Similar patterns have been observed with bacteria that cause brain infections, such as *Neisseria meningitides* and Group B Streptococcus, which adhere in specific locations and flow conditions ^27-29^. To investigate whether mycobacteria exhibit preferential attachment to the brain microvasculature, we use the natural fish pathogen, *Mycobacterium marinum*, which shares 85% of its genes with *M. tuberculosis*^30^. Transgenic zebrafish larvae with green fluorescent vascular endothelial cells (flk:GFP) ^31^ were infected with red fluorescent *M. marinum* and imaged at 1 and 3 days post infection (dpi). At both timepoints, *M. marinum* microcolonies predominantly attached to vessel bifurcations rather than along straight vessels (Figure 1A-D). Given that increased turbulent blood flow at bifurcations is a contributor to aneurysm and stroke ^25,26^, our data suggest that similar hemodynamic forces may influence early interactions between *M. marinum* and brain blood vessels.

**Figure 1:**
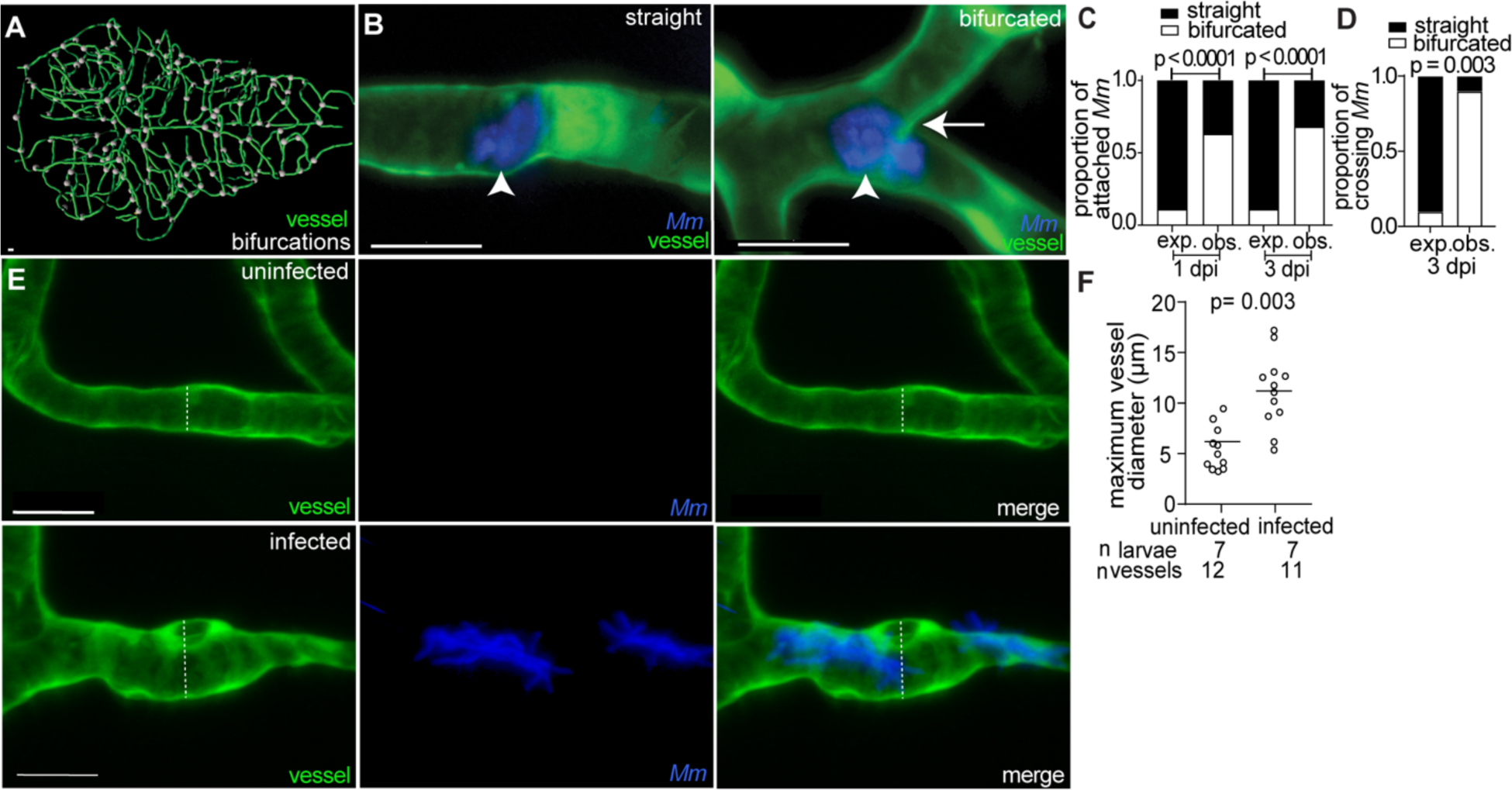
Mycobacteria microcolonies adhere to bifurcations and alter vessel morphology. (A) Rendered trace of brain vasculature of 6 dpf larva. White points denote bifurcations. (B) Representative confocal images of straight (left) and bifurcated (right) vessels from larvae with green fluorescent blood vessels and blue fluorescent *Mm.* Arrowhead, attached microcolony. Arrow, bifurcation. (C,D) Proportion of microcolonies attached (C) and crossing (D) at straight (black) or bifurcated (white) vessels at 1 and 3 dpi. Observed (obs.) proportion is compared to expected (exp.) proportion normalized to total straight or bifurcated vessel length. Fischer’s exact test. (E) Representative confocal images of uninfected (top), and infected (bottom) vessels from larvae with green fluorescent blood vessels and blue fluorescent *Mm* at 3 dpi. Dashed line, diameter of vessel measure in (F). (F) Quantification of maximum diameter of infected and uninfected vessels from the same larvae at 3 dpi. Horizontal bars, means; paired t-test. Scale bar, 10μm throughout.

We next examined vascular morphology following mycobacterial attachment. Cerebral angiography in TBM patients frequently reveals beaded vessels with alternating regions of narrowing and dilation ^5^. To assess whether zebrafish vasculature exhibits similar morphological changes, transgenic larvae expressing green fluorescent endothelial markers (*flk:moesin-GFP*) ^19^ were infected with blue fluorescent *M. marinum* and imaged at 3 dpi. Compared to contralateral uninfected vessels in the same larvae, microcolony-associated vessels displayed significantly increased diameters around bacterial microcolonies (Figure 1E-F), closely resembling the beading morphology observed in TBM patients. These findings indicate that mycobacterial brain infection in zebrafish recapitulates key vascular patterns and morphological features associated with TBM vascular pathology in humans.

### Mycobacterial attachment is sufficient to obstruct blood flow

Infarctions are areas of necrotic tissue resulting from prolonged ischemia and are a hallmark of TBM ^32^. However, the precise mechanism of how they occur in TBM remains unclear ^4^. Thrombosis has been implicated as a potential contributor to infarction early in brain infection ^33-35^, but reports vary on the frequency of organized thrombus formation ^36^. Organized thrombi typically consist of red blood cells (RBCs), platelets, neutrophil extracellular traps, and fibrin ^37^. Once formed, thrombi can occlude blood vessels, restricting oxygen and glucose perfusion and ultimately leading to infarction ^38^.

To investigate the role of thrombosis during mycobacterial brain infection, we intravenously injected blue fluorescent *M. marinum* into transgenic zebrafish larvae with green fluorescent endothelial cells and red fluorescent RBCs (*flk1:GFP;gata1a:DsRed*) ^18,31^. Confocal imaging revealed that RBCs move freely through most uninfected vessels. This is seen as diagonal streaks of red representing RBCs flowing quickly through the vessel during image acquisition (Figure 2A-C). In contrast, most infected vessels contained stagnant RBCs, suggesting occlusion of the vessel (Figure 2A-C). Vessels infected by microcolonies contained immobilized RBCs around the bacteria (Figure 2D-E). However, in other cases vessel occlusion occurred despite the absence of immobilized RBCs (Figure 2D-E). Notably, microcolonies alone were as likely to occlude vessels as those with immobilized RBCs (Figure 2E). Among all infected vessels, blood flow was absent or limited compared to uninfected vessels (Figure 2E), although some uninfected vessels also lacked blood flow (Figure 2C).

**Figure 2:**
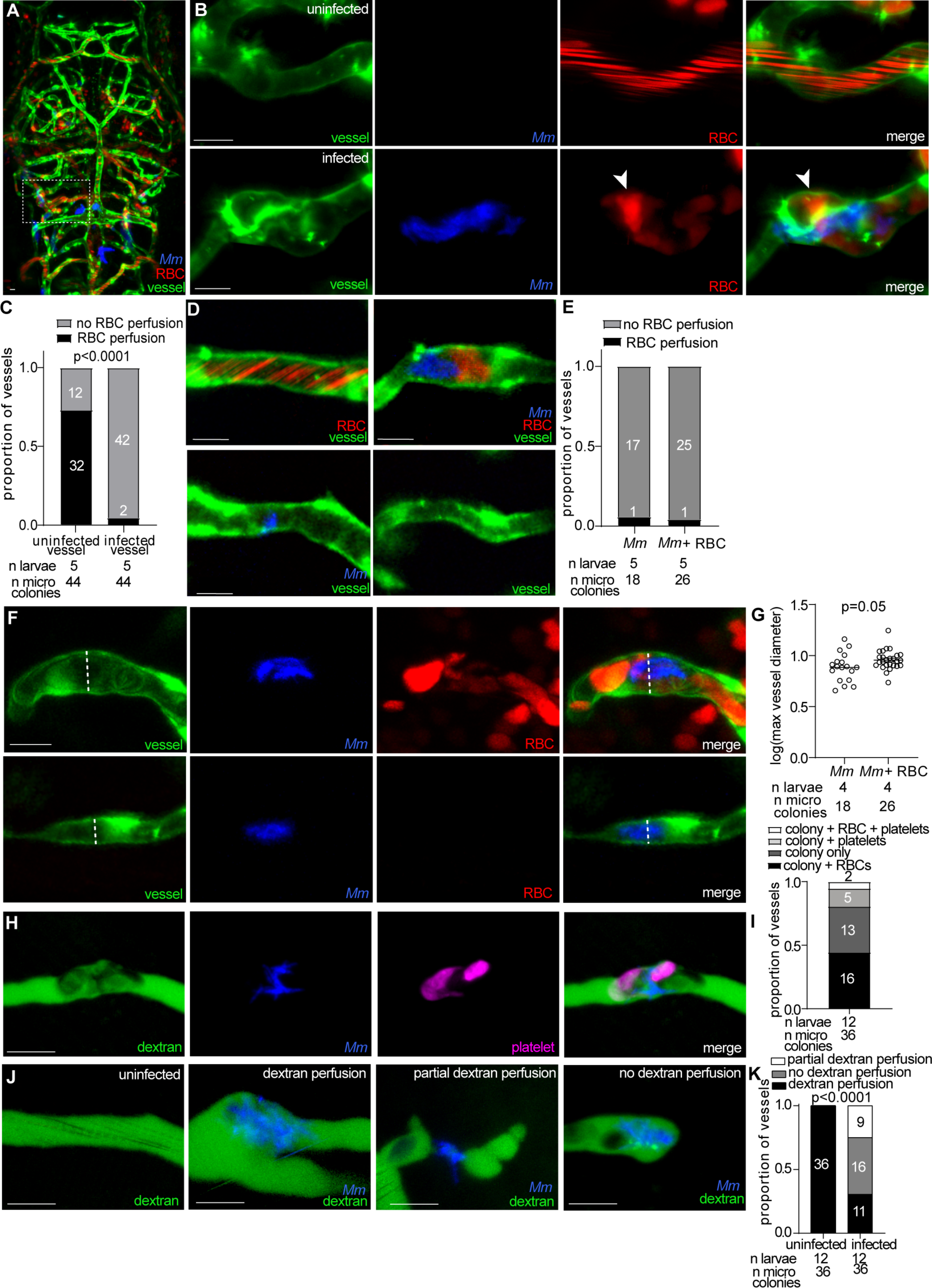
Mycobacteria microcolonies occlude brain blood vessels. (A-B) Representative confocal images of the larva brain vasculature with green fluorescent blood vessels, infected and red fluorescent red blood cells with blue fluorescent *Mm.* Dashed box indicates location of uninfected (top) and infected (bottom) vessels in (B). Arrowhead, stagnant RBCs. (C) Proportion of infected vessels with RBC perfusion in vessels with or without stagnant RBCs around microcolony; Fisher’s exact test. Representative of 2 independent experiments. (D) Brain blood vessels with or without associated microcolonies and/or RBCs. (E) Proportion of infected or uninfected vessels with RBC perfusion; ns: not significant, Fisher’s exact test. Representative of 2 independent experiments. (F) Representative confocal images of larva with green fluorescent blood vessels and red fluorescent RBCs, infected or not with blue fluorescent *Mm.* Dashed line indicates diameter of infected vessel with (top) or without (bottom) stagnant RBCs quantified in (G). (G) Quantification of vessel diameter in infected vessels with or without stagnant RBCs. Horizontal bars, means; Student’s t-test. Representative of 2 independent experiments. (H) Representative confocal images of blue fluorescent *Mm*-infected larva with green fluorescent platelets (pseudo-colored magenta) and intravenously injected Alexafluor647 dextran (pseudo-colored green). (I) Proportion of infected vessels associated with stagnant RBCs and/or platelets. (J) Representative confocal images of blue fluorescent *Mm* infected larva with intravenously injected Alexafluor647 dextran (pseudo-colored green). (K) Proportion of infected or uninfected vessels with no (grey), partial (white) or complete (black) dextran perfusion. Chi-squared test. Scale bar, 10 μm throughout.

Thrombus-associated vessel occlusion often leads to vessel dilation due to increased intraluminal pressure and weakening of the vessel wall ^39^. While infected vessels were generally wider than uninfected vessels (Figure 1E-F), vessels with immobilized RBCs exhibited significantly greater diameters than infected vessels without RBCs (Figure 2F-G), suggesting that RBC accumulation contributes to vascular dilation.

Another mediator of coagulation and thrombosis are platelets ^40^. To examine their involvement in mycobacterial brain infection, we intravenously injected blue fluorescent *M. marinum* into transgenic larvae with green fluorescent platelets and red fluorescent RBCs (*cd41:GFP; gata1a:DsRed*) ^20^. Of vessels with microcolonies, only 19% were associated with platelets, while 6% had both RBCs and platelets (Figure 2H-I), indicating that platelets may play a minor role in vessel occlusion.

In addition to RBC accumulation, vessel occlusion can restrict the passage of luminal contents such as glucose, a critical substrate for neuronal metabolism ^38^. To assess vessel permeability, larvae infected with blue-fluorescent *M. marinum* were injected intravenously with a 10 kDa Alexafluor647 dextran. While all uninfected vessels showed complete dextran perfusion, infected vessels displayed varying degrees of perfusion restriction (Figure 2J). Among infected vessels, 31% maintained full dextran perfusion, 44% exhibited partial perfusion, and 25% were entirely non-perfused (Figure 2J-K). These findings indicate that mycobacterial attachment to brain vessels is sufficient to impair perfusion of luminal contents, independent of thrombus formation.

Together, these data demonstrate that organized thrombi are infrequent in early mycobacterial brain infection and not required for vessel occlusion. Instead, mycobacterial microcolonies alone can obstruct blood flow and limit perfusion of luminal contents throughout the brain.

### Mycobacteria-associated occlusions are transient

Vascular occlusion has been observed in TBM patients, yet its duration remains unclear^41^. To investigate the kinetics of occlusion formation during mycobacterial infection, transgenic larvae with fluorescent vascular endothelial cells and RBCs (*flk1:GFP;gata1a:DsRed*) were infected with blue fluorescent *M. marinum*, and fast-frame imaging was performed for 10 seconds. RBCs remained stagnant in infected vessels (Figure 3A-B), confirming vessel occlusion as observed in still images (Figure 2). In contrast, RBCs in uninfected vessels, from either uninfected or infected larvae, exhibited continuous blood flow (Figure 3A-B). Interestingly, RBC velocity was significantly reduced in uninfected brain vessels from infected larvae compared to those from uninfected larvae (Figure 3A, C), mirroring the hypoperfusion observed in human TBM ^42^.

**Figure 3:**
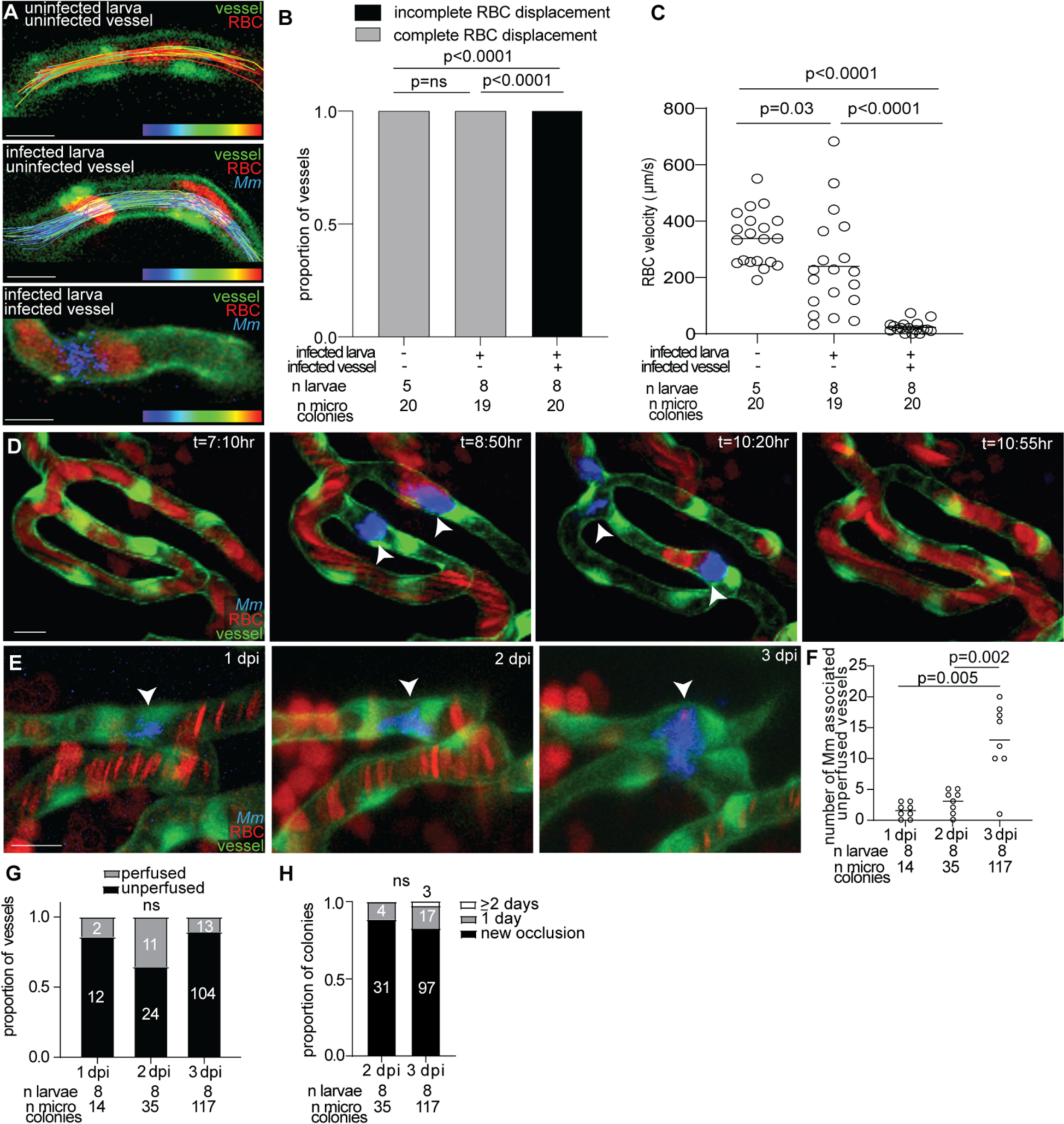
Mycobacteria-associated vessel occlusions are transient. (A) Representative still images from fast frame confocal imaging of larvae with green fluorescent blood vessels and red fluorescent RBCs, infected with blue fluorescent *Mm.* Tracks of individual RBCs are overlain over vessel and color coded from blue (0μm/second) to red (500 μm/second). (B, C) Quantification of RBC displacement (B) and velocity (C) from fast frame imaging. Horizontal bars, means; one-way ANOVA with multiple comparisons. (D) Sequential images from time lapse imaging. Arrowheads, microcolonies obstructing RBC perfusion. t, hours:minutes after start of time-lapse video recording. (E) Sequential images of same microcolony at 1, 2, and 3 dpi. Arrowheads, microcolony obstructing RBC perfusion. Representative of 2 independent experiments. (F) Quantification of number of *Mm* associated occlusions at 1, 2, and 3 dpi. Horizontal bars, means; one-way ANOVA with multiple comparisons. (G) Proportion of infected vessels that are perfused by RBCs at 1, 2, and 3 dpi. ns: not significant; Chi-squared test for trend. (H) Proportion of infected vessels that have a new occlusion (black), or have had an occlusions for 1 (grey) and 2+ days (white). ns: not significant; Chi-squared test for trend. Scale bar, 10 μm throughout.

Time lapse imaging over 14 hours revealed that microcolonies adhered to brain endothelium and transiently blocked blood flow (Figure 3D). All microcolonies detached and reentered circulation within minutes to ∼3 hours of attaching to the vasculature. After detachment, previously occluded vessels regained blood flow (Figure 3D), indicating that early occlusions are transient.

Severe cerebral thrombosis can often last several days to weeks ^32^. To assess longer-term occlusion, larvae were sequentially imaged at 1, 2, and 3 dpi. The number of mycobacteria-associated occlusions increased over time (Figure 3F), though most occlusions lasted for only one day (Figure 3G). These findings indicate that the majority of microcolony-associated occlusions are transient.

### Mycobacteria in vessels is associated with oxidative stress

The ultimate consequence of vascular occlusion is the formation of an infarct, an area of necrotic tissue resulting from prolonged ischemia ^32^. Infarcts have been reported in 13% to 57% of TBM cases based on MRI studies and autopsy findings ^5^. During ischemic stroke, decreased oxygen levels increase the formation of reactive oxygen species (ROS), which drives oxidative stress and leads to cell death and infarct formation ^43^. Given that transient mycobacterial occlusions disrupt blood flow, we sought to determine whether these events induce oxidative stress and subsequent cell death in the brain.

To assess oxidative stress, transgenic zebrafish larvae expressing green fluorescent endothelial cells and red neurons (*flk1:GFP;nbt:DsRed*) ^21^ were intravenously injected with either blue fluorescent *M. marinum* or phosphate buffered saline (PBS) as a control. Prior to imaging, larvae were injected in the hindbrain ventricle with cellROX, a probe that fluoresces when oxidized by ROS. Imaging at 2 dpi, when many microcolonies have adhered to brain blood vessels, revealed significantly higher oxidative stress in infected larvae compared to PBS-injected controls (Figure 4A-B). The oxidative stress was particularly pronounced in both neurons and endothelial cells near vessels containing a microcolony (Figure 4C-D).

**Figure 4:**
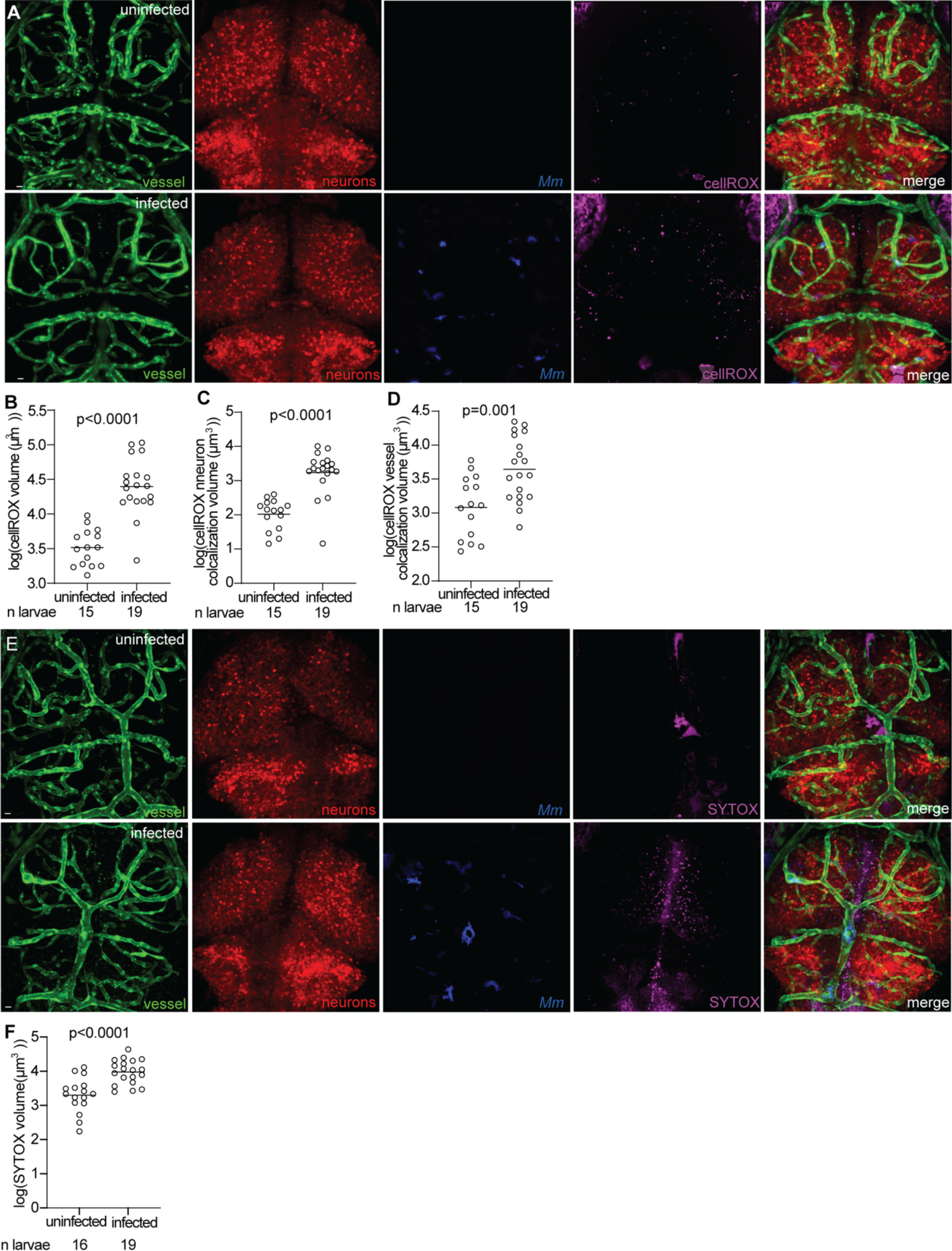
Oxidative stress increases in infected larvae prior to mycobacterial entry into the brain. (A) Representative confocal images of PBS mock injected (top) or blue fluorescent *M. marinum* infected (bottom) 3 dpi larva with green fluorescent blood vessels, red fluorescent neurons, and injected with cellROX. (B) Total cellROX staining volume in infected compared to uninfected brains at 3 dpi. Horizontal bars, means; student’s t-test. (C,D) CellROX staining volume localized to neurons (C) and brain blood vessels (D) in infected compared to uninfected brains at 3 dpi. Horizontal bars, means; student’s t-test. (E) Representative confocal images of PBS mock injected (top) or blue fluorescent *M. marinum* infected (bottom) 3 dpi larva with green fluorescent blood vessels, red fluorescent neurons, and injected with SYTOX. (F) Total SYTOX staining volume in infected compared to uninfected brains at 3 dpi. Horizontal bars, means; student’s t-test. Scale bar, 10 μm throughout.

To determine whether the observed increase in ROS led to cell death, we performed a parallel experiment using SYTOX, which marks dying cells. Transgenic larvae with fluorescent endothelial cells and neurons (*flk1:GFP;nbt:DsRed*) ^21^ were intravenously injected with either *M. marinum* or PBS, followed by hindbrain ventricle injection of SYTOX just before imaging. Compared to PBS-injected larvae, infected larvae exhibited a significantly higher number of SYTOX-positive cells (Figure 4E-F), indicating increased cell death in the brain parenchyma.

Together, these findings demonstrate that mycobacterial attachment to the brain vasculature is associated with increased oxidative stress and neuronal damage. The transient nature of these occlusions suggests that repeated cycles of obstruction and reperfusion may contribute to cumulative oxidative damage, potentially playing a key role in TBM-associated neurovascular pathology.

## Discussion

In characterizing neurovascular events during the early stages of mycobacterial brain infection, we have identified key vascular complications associated with *M. marinum* infection in the zebrafish model. Our findings demonstrate that *M. marinum* preferentially attaches to vessel bifurcations in the brain, leading to both global hypoperfusion and local vessel obstruction. These disruptions contribute to oxidative stress accumulation in both the brain vasculature and neurons. Our study establishes the zebrafish as a valuable model for investigating the vascular complications of mycobacterial brain infection.

Until now, our understanding of TBM-associated vascular pathology has relied on clinical observations^5,11,41^. These studies provide valuable documentation of TBM’s human manifestations and inform diagnostic and treatment strategies. However, clinical observations have inherent limitations. Due to the overlap between TBM symptoms and those of other meningitis, as well as the variability of current culturing techniques, TBM diagnosis is often delayed ^2^. As a result, most of our understanding of TBM-related vascular pathologies pertains to advanced stages of infection or post-mortem findings, leaving the earliest vascular events poorly defined. Additionally, determining causal relationships and molecular mechanisms in human studies is inherently challenging. Animal models provide invaluable opportunities for controlled investigations to uncover causality at the cellular and molecular level. Many studies have demonstrated the utility of zebrafish in modeling TB pathogenesis ^13,22,44,45^. Our findings further support the zebrafish model as a platform for investigating TBM-associated vascular pathology, offering new insights into disease progression.

A debated topic in TBM pathology is the role and prevalence of organized thrombi in vascular occlusions. Human studies have reported conflicting findings regarding the presence of organized thrombi and the duration of vascular events leading to infarction ^5,33-36^. Our study demonstrates that organized thrombus formation is uncommon, and early mycobacteria associated occlusions are predominantly transient. However, we also found occasional thrombus-associated occlusions that persisted for at least 3 days (Figure 3). The presence of multiple occlusion mechanisms in our model may help reconcile discrepancies in human studies, as differences in sampling timepoints and disease progression could account for the variability in thrombus detection.

Vascular complications are a major contributor to long-term disability and death in TBM^6^. However, our current understanding of the onset and progression of these complications remains limited. This knowledge gap hampers the development of effective diagnostic tools and therapeutic interventions ^5^. By extending the use of zebrafish larvae as a model for TBM-associated vascular pathology, our work provides a powerful tool for further mechanistic investigations. Future studies leveraging this model may elucidate novel therapeutic targets, improving clinical outcomes for TBM patients.

## Acknowledgements

The authors thank Richard Daneman for discussion, mentorship, and guidance throughout the project; and Morgan Reitano, Liam Morrill, and the University of California, San Diego aquatics facility staff for zebrafish maintenance.

## Supplemental Materials

Video S1.

Fast frame 83.3 fps confocal imaging of an uninfected green fluorescent blood vessel (*flk1:GFP)* and red fluorescent RBCs (*gata1a:DsRed)* in a PBS-injected larva.

Video S2.

Fast frame 83.3 fps confocal imaging of an uninfected green fluorescent blood vessel (*flk1:GFP)* and red fluorescent RBCs (*gata1a:DsRed)* in a larva infected with blue fluorescent *Mm*.

Video S3.

Fast frame 83.3 fps confocal imaging of an infected green fluorescent blood vessel (*flk1:GFP)* and red fluorescent RBC (*gata1a:DsRed)* in a larva infected with blue fluorescent *Mm*.

Video S4.

Time laspe confocal imaging of a larva with green fluorescent blood vessel and red fluorescent RBCs in a larva infected with blue fluorescent *Mm.* Time lapse images were acquired every 5 minutes for 15 hours.

